# C-terminal tail polyglycylation and polyglutamylation alter microtubule mechanical properties

**DOI:** 10.1101/791194

**Authors:** Kathryn P. Wall, Harold Hart, Thomas Lee, Cynthia Page, Taviare L. Hawkins, Loren Hough

## Abstract

Microtubules are biopolymers that perform diverse cellular functions. The regulation of microtubule behavior occurs in part through post-translational modification of both the *α*- and *β*- subunits of tubulin. One class of modifications is the heterogeneous addition of glycine and glutamate residues to the disordered C-terminal tails of tubulin. Due to their prevalence in stable, high stress cellular structures such as cilia, we sought to determine if these modifications alter the intrinsic stiffness of microtubules. Here we describe the purification and characterization of differentially-modified pools of tubulin from *Tetrahymena thermophila*. We found that glycylation on the *α*-C-terminal tail is a key determinant of microtubule stiffness, but does not affect the number of protofilaments incorporated into microtubules. We measured the dynamics of the tail peptide backbone using nuclear magnetic resonance spectroscopy. We found that the spin-spin relaxation rate (R2) showed a pronounced decreased as a function of distance from the tubulin surface for the *α*-tubulin tail, indicating that the *α*-tubulin tail interacts with the dimer surface. This suggests that the interactions of the *α*-C-terminal tail with the tubulin body contributes to the stiffness of the assembled microtubule, providing insight into the mechanism by which glycylation and glutamylation can alter microtubule mechanical properties.

**SIGNIFICANCE:** Microtubules are regulated in part by post-translational modifications including the heterogeneous addition of glycine and glutamate residues to the C-terminal tails. By producing and characterizing differentially-modified tubulin, this work provides insight into the molecular mechanisms of how these modifications alter intrinsic microtubule properties such as flexibility. These results have broader implications for mechanisms of how ciliary structures are able to function under high stress.

## INTRODUCTION

Microtubules are biopolymers involved in diverse cellular processes within cells including mitosis, cellular transport and signaling, assembly of cilia and flagella for cellular movement, and cellular cytoskeleton structure (1–5). The diversity of function arises in part from post-translational modifications (6, 7). Modifications can occur on both the ordered tubulin body or the disordered C-terminal tails of the *α*- and *β*-tubulin monomers (summarized in Fig. 1). Functionally different subsets of microtubules vary in the degree and type of post-translational modification. For example, acetylation on the interior lumen of microtubules is observed preferentially on stable microtubules (8) and is thought to protect those long-lived microtubules from buckling (9). Microtubule stability is also correlated with detyrosination, which is the removal of the terminal tyrosine on *α*-tubulin (10, 11). Microtubules undergoing dynamic processes, such as mitosis, contain mainly tyrosinated tubulin (12); while, stable and longer-lived microtubules in neurons and cardiomyocytes contain predominantly detyrosinated tubulin (13).

**Figure 1:**
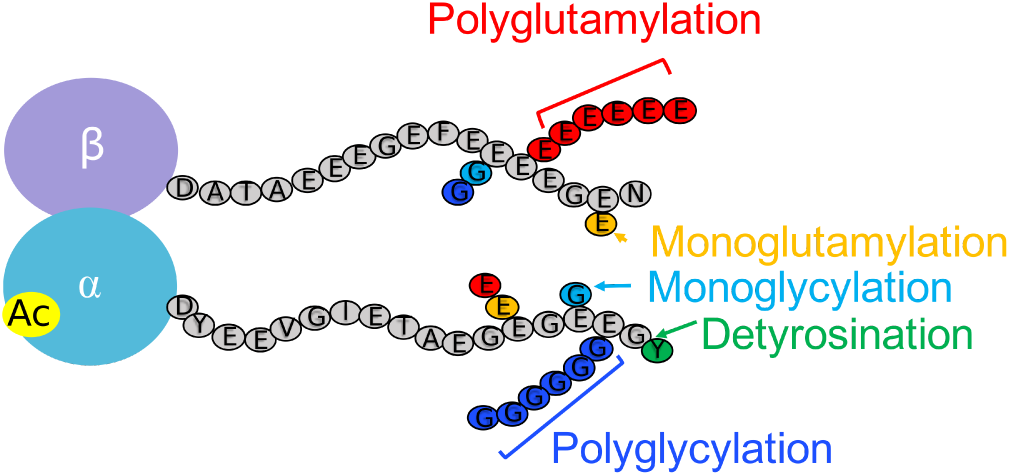
Summary of post-translational modifications on tubulin. Acetylation on *α*-tubulin is located on the ordered tubulin body (yellow). Many modifications are present on the C-terminal tails; the C-terminal tail sequence shown here is that of *T. thermophila*. The terminal tyrosine of *α*- tubulin in many organisms can be removed by detyrosination (green). Both C-terminal tails can contain heterogeneous mixtures of glycylation (blue) and glutamylation (red). These modifications are both mono- and poly-branching additions on the glutamate residues of the C-terminal tails.

Post-translational modifications on tubulin can tune intrinsic microtubule properties, such as flexibility, stability, and polymerization or depolymerization rates. Acetylation increases the mechanical resilience of microtubules, which has been attributed to an increase in the plasticity of the lattice and resulting microtubule flexibility (9). This resilience protects long-lived microtubules from damage (9). Axonemal microtubules have two-fold slower growth and lower catastrophe frequency compared to bovine brain microtubules (14). However, samples with distinct patterns of post-translational modifications prepared by fractionating axonemes show indistinguishable microtubule polymerization rates (14).

A poorly understood class of modifications is heterogeneous additions of glycine and glutamate residues to the C-terminal tails. These modifications result from the stepwise additions to a subset of glutamate residues in the C-terminal tail by tubulin tyrosine ligase-like (TTLL) enzymes (Fig. 1). Glycylation and glutamylation are preferentially found on stable microtubule structures. Glutamylation is present in long, stable microtubules in the axons of neurons (15) and is important for regulation of microtubule turnover and the recruitment of microtubule severing enzymes, such as katanin and spastin (16–18). Glycylation is primarily found on cilia, but the functional role of polyglycylation is not well-understood (19–21). For example, little is known about binding partners specific to this modification. Given the important role of glycylation and glutamylation in cellular structures with particularly high stresses on microtubules (such as cilia), we questioned whether these modifications may directly increase the stiffness of microtubules.

Ciliary microtubules are heavily modified by glycylation and glutamylation. Glycylation is present on the length of mature cilia and may contribute to overall cilia stability (22). In addition, glycylation is present on microtubule structures that anchor the basal bodies to the cell cortex (23). In contrast, glutamylation in cilia is important for early ciliary maturation (24) and proper ciliary beating (25). These two modifications compete for the same sites on the C-terminal tails, and their regulation is clearly coupled (26). The deletion or depletion of glycine ligases in cells results in a compensatory increase in glutamylation (27); similarly, mutations of the modification sites on one of the tubulin subunits affect the levels of glycylation and glutamylation on the opposing non-mutated C-terminal tail (24, 28). The perturbation of the balance of these modifications causes shorter cilia (27), destabilization of already assembled cilia (26), and reduced frequency of primary cilia (29). While extrinsic changes in protein interactions with microtubules could cause these cellular effects (30), we hypothesize that there are intrinsic alterations in stiffness due to post-translational modifications.

Although they comprise only a small percent of the total tubulin mass and make only transient interactions with adjacent dimers, the C-terminal tails appear to alter microtubule flexibility. Changes in divalent salts can change microtubule flexibility in a C-terminal tail dependent manner (31), indicating that tails interact sufficiently with either each other or the surface of the tubulin body to alter microtubule flexibility. Simulations indicate that the C-terminal tails are sufficiently collapsed such that they may not interact with each other. However, simulations do show the interaction of the *α*-tubulin tail with clusters of positive residues on the *β*-tubulin body surface on the neighboring dimer and interactions of the *β*-tubulin tail with a cluster of positive amino acids on the adjacent *α*-tubulin surface (32). One proposed difference between glycylation and glutamylation is the intrinsic rigidity of the modified polypeptide; for example, glycylated C-terminal tails may be more collapsed that glutamylated ones because of their relatively higher hydrophobicity and lower electrostatic repulsion (33).

Nuclear magnetic resonance spectroscopy (NMR) is an ideal tool for determining the atomic level behavior of the C-terminal tails and can be used to detect even transient interactions. It is the primary structural tool for intrinsically disordered proteins, which are refractory to electron microscopy and x-ray crystallography approaches. In NMR experiments, the C-terminal tails of glycylated tubulin were observed in two states, both different than isolated peptides. This suggests that the conformation of the C-terminal tails changes as a result of interactions with the surface of the tubulin body (34).

To address whether and how glycylation and glutamylation affect the intrinsic flexibility of microtubules requires samples of differentially-modified tubulin. Current purification of porcine brain tubulin yields tubulin containing a mixture of modifications. Two approaches to purify differentially modified tubulins have been developed. In the first, recombinant unmodified human tubulin is purified and subsequently modified using purified glycine and glutamate ligases (members of the TTLL family of enzymes) (35). In contrast, Souphron et al. used genetic means to control the level of modification in the tubulin source, as we do here (36). These existing purification strategies are not suitable for the production of isotopically-enriched protein for the purposes of structural investigation of the C-terminal tails by NMR. Our method of purifying tubulin from *T. thermophila* gives the unique ability to address the molecular behavior of the C-terminal tails and how they may affect the flexibility of microtubules (34).

In this study, we purified differentially-modified tubulin from different *T. thermophila* strains and found that microtubules with glycylated *α*-tubulin tails are stiffer than their unmodified or glutamylated counterparts. We used electron microscopy to determine that changes in rigidity are not correlated with changes in protofilament number. On the other hand, we found differences in NMR ^15^N-relaxation properties of *α*- and *β*-C-terminal tails, consistent with the difference in sensitivity to the modifications on the two tubulin tails. As measured by R_2_ spin relaxation, the interactions of the *α*-C-terminal tail residues depend on the distance from the microtubule surface more prominently than do those of the *β*- tubulin tails, suggesting more significant surface interactions.

## MATERIALS AND METHODS

### Growth and purification of tubulin from mutant *T. thermophila* strains

The *Tetrahymena thermophila* strains containing modifications to the C-terminal tail sequence (ATU1-6D, (24)) and knock-outs of the TTLL-3 family of enzymes (TTLL3(A-F)-KO, (27)) were generously gifted by Jacek Gaertig (University of Georgia). *T. thermophila* strains, TTLL3(A-F)-KO and ATU1-6D, were cultured in SPP rich media. The cultures were grown in a shaking incubator at 30°C at 100rpm in 2.8L Fernbach flasks. The cells were harvested at maximum density (1×10^6^ cells/mL) by two 5 minute centrifugations at 2800 x g. Cell pellets were frozen at −70°C until further purification. Live cell movies were taken using Nikon Widefield with a 4X objective in a 100*µ*m well on a glass coverslip. The frame rate was 0.118 sec/frame with an exposure time of 354*µ*s. Tubulin was purified from each strain as previously described. A modified TOG-affinity column was made by published methods as described previously (37). Tubulin purification proceeded using an affinity-based purification on the TOG-affinity column as described previously (34).

### Western blot of tubulin from *T. thermophila* strains

10% SDS-PAGE gels containing 0.5*µ*g of tubulin purified from each of the strains were transferred to a PDVF membrane. Antibodies and respective concentrations were as previously published (27). The primary antibodies and dilutions used were as follows: (1) Mouse Anti-alpha tubulin (Developmental Studies Hybridoma Bank clone 12G10) at 1:250, (2) Rabbit Anti-polyglutamylation (1:5000) (a generous gift from Jacek Gaertig, (24)), (3) Mouse Anti-monoglycylated tubulin (clone TAP 952, EMD Millipore) at 1:5000, (4) Mouse Anti-pan polyglycylated tubulin (clone AXO 49, EMD Millipore) at 1:5000. Secondary antibodies were used at a dilution of 1:7500, and were the following: (1) Anti-mouse IgG (H+L) AP Conjugate (Promega), and (2) Anti-rabbit IgG (Fc) AP Conjugate (Promega). Western blots were developed using the Pierce 1-Step NBT-BCIP reagent as recommended by the manufacturer and imaged with a Canon LiDE 210 Scanner.

### Mass spectrometry of tubulin from *T. thermophila*

Purified tubulin samples were reduced, alkylated, and digested with trypsin (*α*-tail) or chymotrypsin (*β*-tail). Samples were resolved using a Waters nanoAcquity UPLC. Chromatographic separation was performed with a BEH C18 reversed-phase column (250 mm × 75 *µ*m, 1.7 *µ*m, 130 Å, Waters), using a linear gradient from 95% Buffer A (0.1% formic acid) to 35% Buffer B (0.1% formic acid, 99.9% acetonitrile) over 60 min at a flow rate of 300 nl/min. MS/MS was performed using an LTQ-orbitrap mass spectrometer, scanning between 150 to 2,000 m/z (60,000 resolution) and select the top six most intense precursor ions were for MS/MS sequencing using monoisotopic precursor selection, rejecting singly charged ions. Dynamic exclusion was used with a repeat count of one, a repeat duration of 30 s, an exclusion duration of 180 s, and an exclusion mass width of 20 ppm. The maximum injection time for Orbitrap parent scans was 500 ms with a target AGC (automatic gain control) of 1 × 10^6^. The maximum injection time for LTQ MS/MS scans was 250 ms with a target AGC of 1 × 10^4^. The normalized collision energy was 35% with an activation Q of 0.25 for 30 ms.

### Microtubule Assembly

Microtubules were assembled in PIPES-based BRB80 (80mM PIPES pH 6.7, 1mM MgCl_2_, 1mM EGTA) supplemented with 2mM GTP (Sigma Aldrich) at 37°C for one hour, then incubated at room temperature for at least 30 minutes. Starting tubulin concentration ranged from 1-10*µ*M. Samples for fluorescent bending measurements were supplemented with 10% rhodamine-labeled porcine brain tubulin (Cytoskeleton, Denver, CO). The resulting microtubules were spun down at 16300 x g in a table-top microcentrifuge. Pelleted microtubules were resuspended in BRB80 supplemented with 10*µ*M taxol. High magnesium conditions were supplemented with 50mM MgCl_2_.

### Electron Microscopy of Microtubules

After assembly, the microtubules were spun at 16300 x g in a table-top microcentrifuge. The microtubules were resuspended in BRB80 supplemented with 10*µ*M taxol. Approximately 4*µ*l of undiluted or diluted (1:10) microtubules were adsorbed to glow discharged holey-carbon C-flat grids (Protochips, Inc, Raleigh, NC) for 10 to 30 seconds, blotted with Whatman filter paper, and immediately plunge-frozen into liquid ethane using a homemade plunge-freezing device. Frozen samples were transferred under liquid nitrogen to a Gatan-626 cryo-holder (Gatan, Inc, Pleasanton, CA). Cryoelectron microscopy data was collected on an FEI Tecnai F20 FEG transmission electron microscope (FEI-Company, Eindhoven, The Netherlands) operating at 200kV. Images were collected at a magnification of 29,000x and a defocus of −4.0 *µ*m using a total dose of 33 electrons/Å^2^. Images from wild-type microtubules were recorded binned by two on a 4K × 4K Gatan Ultrascan 895 CCD camera (Gatan, Inc). With this camera at a microscope magnification of 29,000x, the resulting pixel size corresponds to 7.6Åon the specimen. Images from the TTLL3(A-F)-KO microtubules were taken on the same microscope which was fitted with a 24 megapixel, 5.7K × 4.1K Gatan K3 camera (Gatan, Inc). Images were collected at a microscope magnification of 11,500x and a defocus of −3.5 *µ*m using a total dose of 38 electrons/Å^2^. With this camera the images were recorded binned by two with resulting pixel size on the specimen of 6.2Å. Each image was captured as 50 frame movies (0.76 electrons/Å^2^/frame) and aligned using SerialEM software (38). This software was also used to automate the data acquisition and minimize exposure of the specimen to the electron beam.

### Fluorescence Microscopy

All microtubules were assembled in BRB80, as described above. Microtubules were diluted at least 20 fold for imaging in the relevant buffer, with the addition of an oxygen scavenging system (0.5% *β*-mercaptoethanol, 4.5mg/mL glucose, 0.2mg/mL glucose oxidase, and 0.035mg/mL catalase), 10*µ*M Taxol to stabilize the microtubules (Sigma Paclitaxel), and 1% Pluronic F-127 to inhibit sticking to the slide or coverslip (Sigma). 0.25*µ*L of the dilution was sealed with epoxy between a glass coverslip and glass slide and kept in the dark unless being imaged. Slides and coverslips were cleaned in a bath sonicator in 100% ethanol for 45 minutes and soaked in 100% ethanol until use. The slides and coverslips were flamed to dry immediately prior to use. Fluorescence bending images were acquired using a Nikon Confocal Spinning Disc with a 1.45 NA 100X oil objective (Nikon) and Andor 888 Ultra EMCCD camera. Movies were acquired for at least 100 frames. There was a 1 second delay between frames with a 100ms exposure time.

### Analysis of Flexural Rigidity

All movies were cropped to contain a single microtubule using ImageJ (39). Less than ten percent of the total number of frames were removed if the microtubule drifted out of the z-plane or if any other particles obstructed the microtubule during imaging. Single microtubules were analyzed by previous methods (40–42). In short, the variances in the magnitude of first twenty-five normal modes was measured using a customized MATLAB script. To determine the uncertainty in the measured variance, bootstrapping statics was employed using R (43) as previously described (40). Average persistence lengths were determined by fits to the cumulative distribution function (Supplemental Figure S4, Supplemental Table S1).

### ^15^N Relaxation NMR (R_1_/R_2_)

Tubulin was purified from wild-type *T. thermophila* grown in minimal bacterized media as previously described for isotopic labeling (34). GST-CTT peptides were expressed and purified as previously described (34). T_1_ and T_2_ relaxation experiments were performed using the standard ^15^N-HSQC experiment from the Varian BioPac (gNhsqc) on an 800 MHz magnet. For measurement of T_1_, relaxation delays used were: 0, 0.1, 0.2, 0.3, 0.4, 0.5, 0.6, 0.7, 0.8, 0.9 ms. For the measurement of T_2_, relaxation delays used were: 0.01, 0.03, 0.05, 0.07, 0.09, 0.11, 0.13, 0.15, 0.17, 0.19, 0.21, 0.23, 0.25 ms. The data was processed using standard scripts in NMRPipe, and analyzed using CCPNmr Analysis software (44).

## RESULTS AND DISCUSSION

### Creation and characterization of differentially-modified pools of tubulin

To obtain samples with different modifications, we took advantage of existing *T. thermophila* strains created to study the biological role of C-terminal tail modifications (24, 27, 45). We purified tubulin from three strains, giving primarily glycylated tubulin (Wild-type), primarily glutamylated tubulin (TTLL3(A-F)-KO), and tubulin with glycylation only on the *β*-tubulin C-terminal tail (ATU1-6D). As we published previously, tubulin purified from our wild-type strain is primarily glycylated (34).

To obtain glutamylated tubulin, we purified tubulin from a strain with all members of the TTLL3 family of enzymes deleted (27). We refer to this strain as TTLL3(A-F)-KO. Strains lacking the glycine ligases showed increased levels of polyglutamylation (27). This strain has shorter cilia, slower growth rates, and an increased resistance to stabilization by paclitaxel (27). We observed a disrupted swimming pattern (Fig. 2), most likely due to changes in the ciliary length and organization.

**Figure 2:**
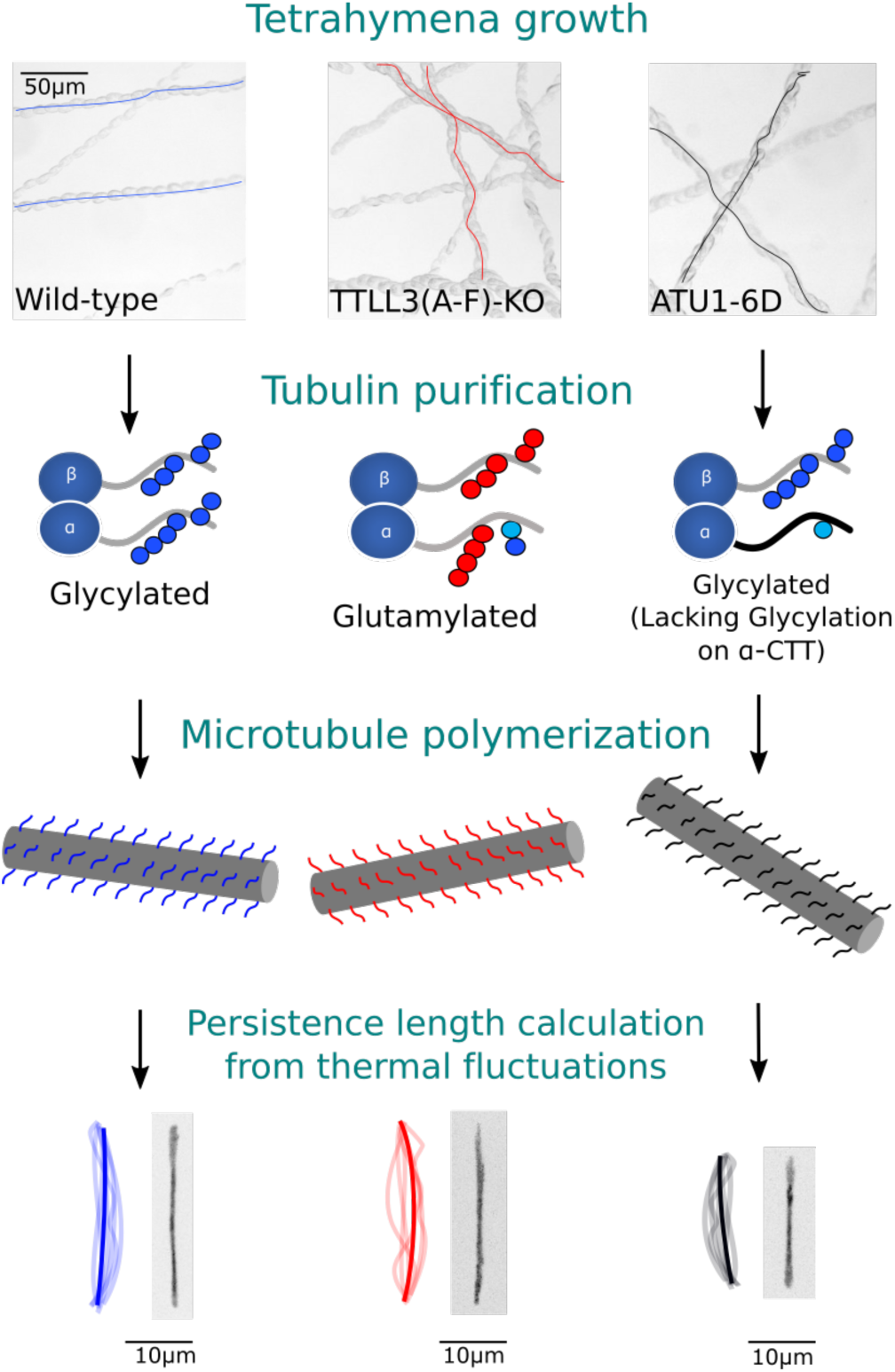
Preparation and analysis of flexural rigidity of differentially-modified microtubules. We purified tubulin from three different strains of *T. thermophila*. The two mutant strains have ciliary defects and abnormal swimming patterns. We characterized the tubulin post-translational modifications by mass spectrometry and western blotting and determined that our samples were primarily glycylated (wild type), glutamylated (TTLL3(A-F)-KO), or glycylated only on the *β*-tubulin tail (ATU1-6D). We then measured the bending rigidity of these different microtubule pools.

To obtain tubulin that was polyglycylated on only *β*- tubulin, we purified tubulin from a strain where the last six glutamate residues of *α*-tubulin were mutated to aspartate (ATU1-6D) (24). We previously mapped the sites of glycylation to these residues, and did not observe glycylation on the other glutamate residues in the *α*-tubulin C-terminal tail (34). The aspartate substitutions retain the overall charge of the tail but disrupt the chemistry of modifications. This strain has slower swimming speeds (24) and disrupted swimming patterns (Fig. 2), though they are able to grow well enough for the purposes of purification of tubulin.

We first used measured levels of glutamylation by western blot using an antibody specific to polyglutamylation. We detected polyglutamylation in tubulin purified from all three strains tested by western blot (Supplementary Materials Fig. S1), including the wild-type strain where glutamlylation was not previously observed by either mass spectrometry or NMR (34). We observed that the tubulin purified from TTLL3(A-F)-KO has significantly higher levels of polyglutamylation on both tails than the other two strains, which is as expected due to previous observations of hyperglutamylation in whole cell lysates of TTLL3(A-F)-KO cells (21, 27). While glycine and glutamate modifications are chemically different, cells appear to compensate for a loss of glycine ligase activity with an increase in compensatory glutamate ligase activity. We detected little polyglutamylation on *α*-tubulin purified from the ATU1-6D strain, as expected because all modifications sites were mutated. However, the levels of polyglutamylation detected on *β*-tubulin from ATU1-6D cells is similar to the wild-type. Glutamylated peptides were diffi cult to detect by mass spectrometry, but were observed in samples purified from TTLL3(A-F)-KO cells.

We then measured the degree of glycylation using both western blot and mass spectrometry. We used antibodies to monoglycylated tubulin (TAP952) and polyglycylated tubulin (AXO49) to be specific to mono or polyglycylation respectively (Supplementary Materials Fig. S1). As previously determined by mass spectrometry, the tubulin purified from wild-type *T. thermophila* contained both mono- and polyglycylation (34). Both tails on *α*- and *β*-tubulin have significant glycylation. Because modification levels are thought to be different on the cilia relative to the rest of the cell, we deciliated cells and purified tubulin from the ciliary and cell body fractions. Unexpectedly, there were no significant differences in mass spectrometry results between tubulin purified from the cell body and shed cilia by mass spectrometry (Fig. 3A,B).

**Figure 3:**
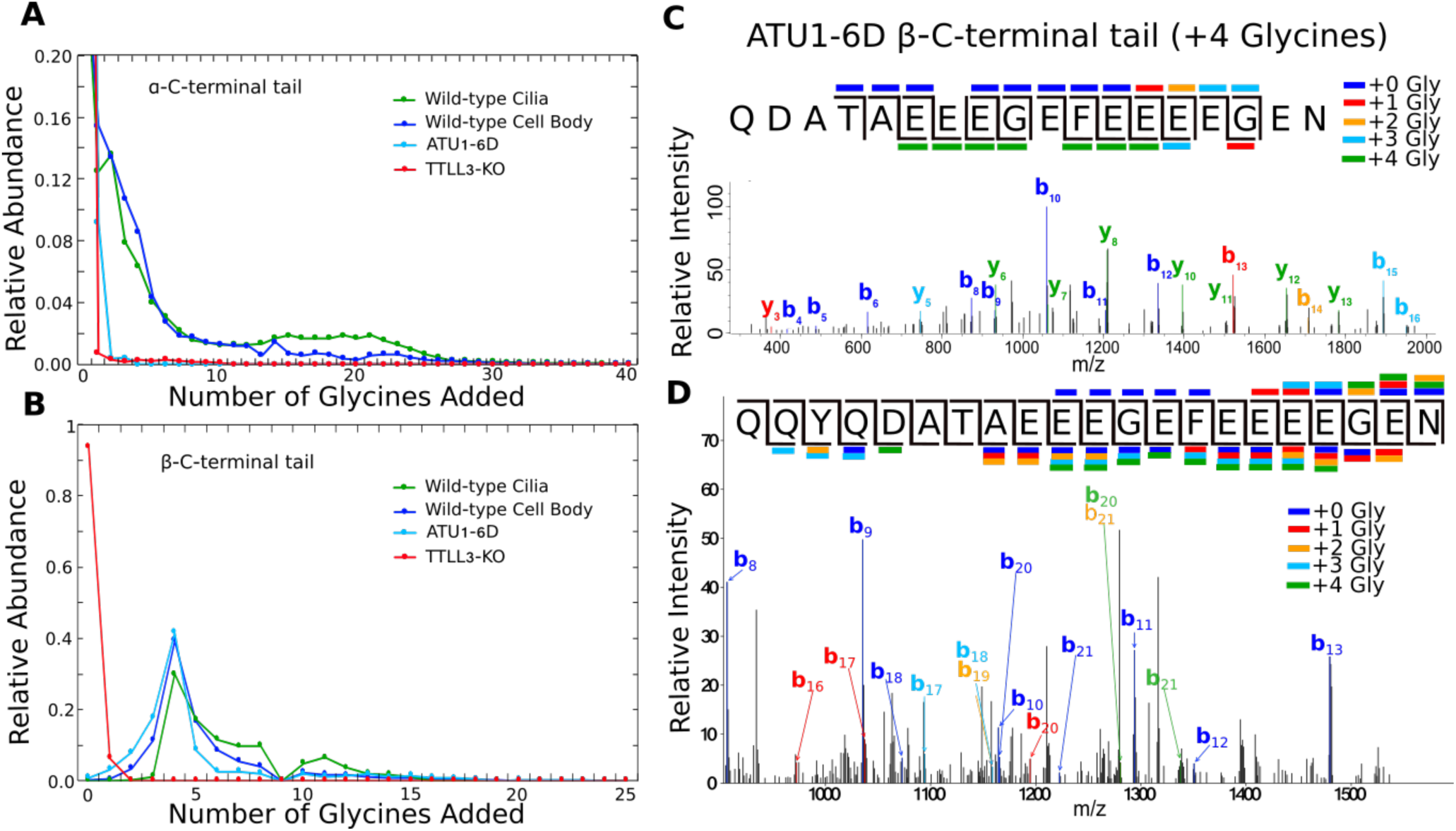
Mass spectrometry characterization of the glyclation patterns on differentially-modified tubulin samples. For the wild-type samples, we purified tubulin from both ciliary and cell body fractions. (A, B) We measured the total ion current corresponding to each C-terminal tail peptide for each of our samples, finding a peak in the *β*-tubulin tail sample of four additional glycines. We mapped the glycyine additions of this parent peptide using MS-MS. (C) An example MS-MS scan showing fragmentation consistent with a single glycine on each of the four terminal glutamates. (D) A different MS-MS scan containing fragments corresponding to a wide rage of glycine localizations. Fragmentation of the modification itself is thought to contribute to the diversity of y-ions, as we did not observe b-ions consistent with modifications upstream of the four terminal glutamates.

We detected no mono- or polyglycylation in tubulin purified by the same protocol from the TTLL3(A-F)-KO strain by western blot. This is as expected because the enzymes responsible for the ligation of glycine residues were deleted from the micronucleus (27). However, by mass spectrometry, we detected peptides corresponding to mono- and polyglycylation on the *α*-C-terminal tail. The low levels of glycylation are presumed to be from residual DNA coding for the TTLL3 enzymes in the macronucleus or could potentially be due to another glycylating enzyme. TTLL3(A-F)-KO tubulin has no detectable polyglycylation on the *β*-C-terminal tail by mass spectrometry.

For the tubulin purified out of the ATU1-6D strain, levels of mono- and polyglycylation on the *β*-tubulin subunit were very similar to that seen in our wild-type cells. Although all glutamate residues that we had previously determined to be sites of modification on the *α*-tubulin tail were mutated in this strain to aspartates, we observed low levels of monoglycylation on the *α*-tubulin subunit by western blot (Supplementary Materials Fig. S1). By mass spectrometry, we detected monoglycylation on the introduced aspartate residues, and not on any residues further upstream on the *α*-C-terminal tail (data not shown). Therefore, the TTLL3 glycine ligases appear to be able to modify the aspartate residues, albeit at a very low level. Interestingly, we detected tyrosinated ATU1-6D *α*-tubulin in addition to detyrosinated ATU1-6D *α*-tubulin; we have not detected the peptide containing the final tyrosine residues in any of our other samples.

We measured the total ion current corresponding to the mass addition of varying numbers of glycine residues for all three strains and found a strong preference for the addition of 4 glycine residues to the *β*-tubulin C-terminal tail in wild type and ATU1-6D cells (Fig. 3A,B). We then analyzed the distribution of glycine residues on this species. The fragmentation pattern in MS-MS experiments of the peptides with four added glycine residues was very heterogeneous. (Fig. 3c,d, Supplementary Materials Fig. S2). We obtained strong evidence for several different arrangements of the four glycine residues. For example, we were able to identify fragments that strongly support the presence of a species with one glycine on each of the last four glutamates from the peptide from ATU1-6D (Fig. 3C). We also identified a MS/MS fragments corresponding to several other heterogeneous arrangements in this ATU1-6D tubulin peptide (Fig. 3C), including all four glycine residues being on the last glutamate residue. However, the fragmentation of not just the tail peptide but also the modification itself confounded our analysis. Although we found y-ions giving evidence of fewer than four additional glycines in the MS-MS fragments, we did not find b-ions containing positive evidence of glycine modifications upstream of the final five glutamates. We conclude the the primary sites of modification are the five C-terminal glutamates, which is similar to the previous mapping of wild-type tubulin (34).

From both the western blot and mass spectrometry analysis, we conclude that we have three distinct subsets of differentially modified tubulin: (1) polyglycylated tubulin from wild-type *T. thermophila*, (2) polyglutamylated tubulin from TTLL3(A-F)-KO *T. thermophila*, and (3) tubulin polyglycylated on only the *β*-tubulin C-terminal tail, with an unmodified *α*-C-terminal tail from ATU1-6D *T. thermophila*. These modifications are summarized in Fig. 2. With these tools in place, we sought to determine the role of glutamylation and glycylation on microtubule mechanical properties.

### Microtubule flexibility depends on the post-translational modifications present on the C-terminal tails

Glycylation and glutamylation are located in microtubule structures where mechanical integrity is important. Cilia exert high forces (10s of pN), and the microtubule stiffness is thought to dominate the cilia mechanical properties (46). Microtubule post-translational modifications are important for cilia length regulation and mechanical integrity. Cells with reduced levels of either glutamylation or glyclation have shorter cilia and abnormal swimming (24, 27). We hypothesized that the modification state of ciliary microtubules may affect their overall stiffness.

In order to test the hypothesis that microtubule stiffness varies between differentially-modified microtubules, we measured the persistence length (L_*p*_) of each freely fluctuating microtubules from our three tubulin pools using fluorescence microscopy as was done previously (40–42, 47). We analyzed the distribution of contributions of each normal modes to the microtubule shape (47). We acquired frames every second for eight minutes. The frames were approximately uncorrelated. We used bootstrapping to ensure good error estimation and for robustness of the analysis (40). From these measurements, we determined the microtubule persistence length, which is a measure of the stiffness of a filament. A stiffer filament has a higher persistence length, while a lower persistence length means a more flexible filament. Consistent with previous work, we found no dependence of persistence length on contour length (40, 42, 47, 48), which ranged from 10*µ*M to 45*µ*M (Fig. 4, Supplementary Materials Fig. S3). We fit the resulting cumulative distribution functions to determine the average persistence length for each tubulin pool (41).

**Figure 4:**
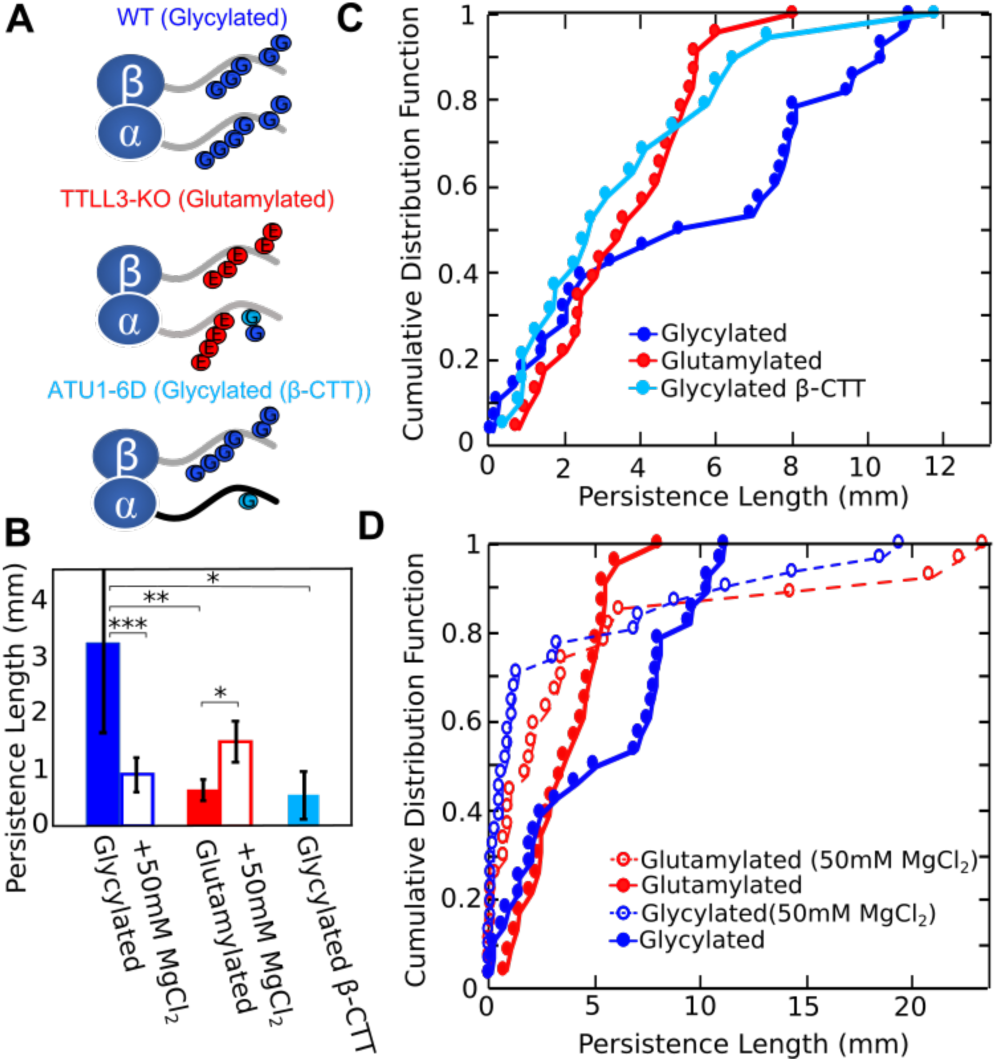
Measured persistence lengths of our differentially-modified microtubules. (A) A summary of the modification states as determined. (B) The average and standard error of the measured persistence lengths. (C, D) Cumulative distribution functions for our different microtubule populations. Pairwise p-values are listed in Supplementary Material Table S1, and all pairs with p-value of p<0.05 (*), p<0.01 (**), and p<0.001 (***) by the K-S test are indicated.

We tested whether differences in C-terminal tail post-translational modifications, in particular glycylation and glutamylation, affected the stiffness of microtubules by comparing the persistence lengths of microtubules made from our three tubulin pools (Fig. 4). Microtubules from *T. thermophila* tubulin ranged in persistence length from L_*p*_ = 0.64 ± 0.18 mm for glutamalyted (n=23) microtubules to L_*p*_ = 3.24 ± 1.60 mm for polyglycylated microtubules (n=28). The increased stiffness of polyglycylated *T. thermophila* microtubules may contribute to the high stability of cilia.

Because not all of the measurement sets were normally distributed, we use the non-parametric Kolmogorov-Smirnov statistical test to determine the statistical likelihood of different measured persistence lengths differed (49). The pairwise p-values for all of our data are listed in Supplementary Materials (Table S1). The average persistence length of tubulin purified from the glycylated microtubules (wild type cells, L_*p*_ = 3.24 ± 1.60 mm; n=28) was five-times stiffer than that of tubulin from either glutamylated (TTLL3(A-F)-KO cells, L_*p*_ = 0.64 ± 0.18 mm; n=23) or glycylation on the *β*-C-terminal tail only (ATU1-6D cells, L_*p*_ = 0.55 ±0.42 mm; n=19), and these distributions showed strong statistical significance (p-value = 0.007 and p-value = 0.042, respectively). Cumulative distribution functions of the persistence length measurements are shown in Fig. 4C. The persistence length measurements of TTLL3(A-F)-KO tubulin were very similar to those of ATU1-6D tubulin. This is surprising, as our mass spectrometry results indicate that ATU1-6D *β*-tubulin tails contain similar post translational modifications to that of the wide type tubulin, but are distinctly different than that of the TTLL3(A-F)-KO tubulin. In contrast, the *α*-tubulin tails differ significantly in all three cases.

Our results suggest that glycylation on the *α*-C-terminal tail is a key determinant of the stiffness of microtubules. Removal of glycylation on only the *α*-C-terminal tail (while retaining glycylation on the *β*-C-terminal tail) decreases the stiffness of microtubules compared to microtubules containing glycylation on both tails. Microtubules assembled from tubulin with glycylation on the *β*-C-terminal tail only (ATU1-6D) have a similar distribution of persistence length as the glutamylated microtubules (TTLL3(A-F)-KO). This observation suggests that glycylation on the *β*-C-terminal tail alone is not sufficient to increase the stiffness of microtubules, supporting a biological difference in function between the modifications on the two tails.

Polyglycylated microtubules are approximately three-fold stiffer than taxol-stabilized rhodamine-labeled porcine tubulin (1.19 ± 0.04 mm) (40–42). In contrast the polyglutamylated *T. thermophila* microtubules (0.64 ± 0.18 mm and n=23) are slightly more flexible than porcine tubulin. The differences between porcine and *T. thermophila* tubulin could be attributed to sequence and structural differences between the two species of tubulin. In addition, porcine brain tubulin contains a different mixture of post-translational modifications than do our samples, including potentially acetylation which is known to reduce microtubule stiffness (8, 9). However, our samples do not have significantly less acetylation than reported values for bovine brain tubulin (∼30% acetylated) (50). We compared the ratio of ion-currents (semiquantitative analysis) to determine the level of acetylation in our samples and found the level of acetylation in our wild type sample to be 50%, our ATU1-6D sample to be about 16%, and our TTLL3(A-F)-KO at 37%. There does not appear to be strong correlation between stiffness and acetylation level in our samples.

We sought to determine whether the divalent cation magnesium would affect bending rigidity differently for our glycylated and glutamylated microtubules. Divalent salts have been shown to affect microtubule bending rigidity in a C-terminal tail dependent manner (31). The high negative charge of the tails is thought to be important for their contribution to the salt dependence. Microtubule glutamylation adds significant charge to the C-terminal tails, while glycylation does not, and may instead act as a steric barrier to prevent charges from approaching each other. Furthermore, the conformation of the C-terminal tails may depend on the divalent salt concentration, and may affect the flexibility of microtubules (31). We performed similar measurements as above of persistence length of glycylated versus glutamylated microtubules in very high magnesium conditions (50mM MgCl_2_).

We observed significant changes in the measured persistence lengths between buffers containing low or high magnesium concentrations (1 mM or 50 mM MgCl_2_ respectively) (Fig. 4d). However, we found no significant difference in the distribution of persistence lengths for the two microtubule subsets (glycylated or glutamylated) relative to each other in the high magnesium condition. Interestingly, a small subset of both glycylated and glutamylated microtubules had persistence lengths of greater than 10mm with the added magnesium (Fig. 4). We could not identify a reason for these stiffer filaments; they did not appear to be bundled or to incorporate a different concentration of fluorescently labeled tubulin. Contrary to our expectation, the glycylated microtubules showed more significant changes in bending rigidity than did the glutamylated ones. The glycylated microtubules are significantly more flexible in high magnesium than in low magnesium conditions. However, the flexibility of the glutamylated microtubules under high magnesium conditions did not significantly change. Glycylated microtubules appear more sensitive to changes in the external salt conditions, specifically divalent salts.

Our results are in contrast to previous work on mammalian tubulin where the addition of divalent salts increased the persistence lengths of microtubules (31). However, glycylation is not a predominant modification in porcine tubulin, indicating that the effect of divalent cations might be dependent on the modifications present on the C-terminal tails. The differences in the effect of glycylation and glutamylation on microtubule stiffness may be due to the conformation of the C-terminal tails on the surface. Glycylated tails are hypothesized to adopt a collapsed conformation (33). The higher salt in the surrounding environment may disfavor this conformation and promote a conformation similar to the glutamylated microtubules.

### Differentially-modified microtubules incorporate similar numbers of protofilaments

We next tested whether differences in bending rigidity of the different microtubule pools could be explained by structural differences in the assembled microtubules. In mammalian tubulin, microtubules containing different numbers of protofilaments showed different microtubule flexural rigidity (41). We used electron microscopy to determine if tubulin modifications caused changes in the number of protofilaments incorporated into microtubules. We considered only the wild-type and TTLL3(A-F)-KO strains, which are primarily glycylated and glutamylated respectively. The number of protofilaments per microtubule is typically 13 in cells, but can vary from 9-17 under *in vitro* polymerization conditions (51). In electron cryo-microscopy images, the number of protofilaments in each microtubule can be determined from the unique helical pitch and moire patterns of each microtubule type (51). We counted the number of protofilaments for wild-type (glycylated) and TTLL3(A-F)-KO (glutamylated), n=145 and n=96 microtubules respectively (Fig. 5).

**Figure 5:**
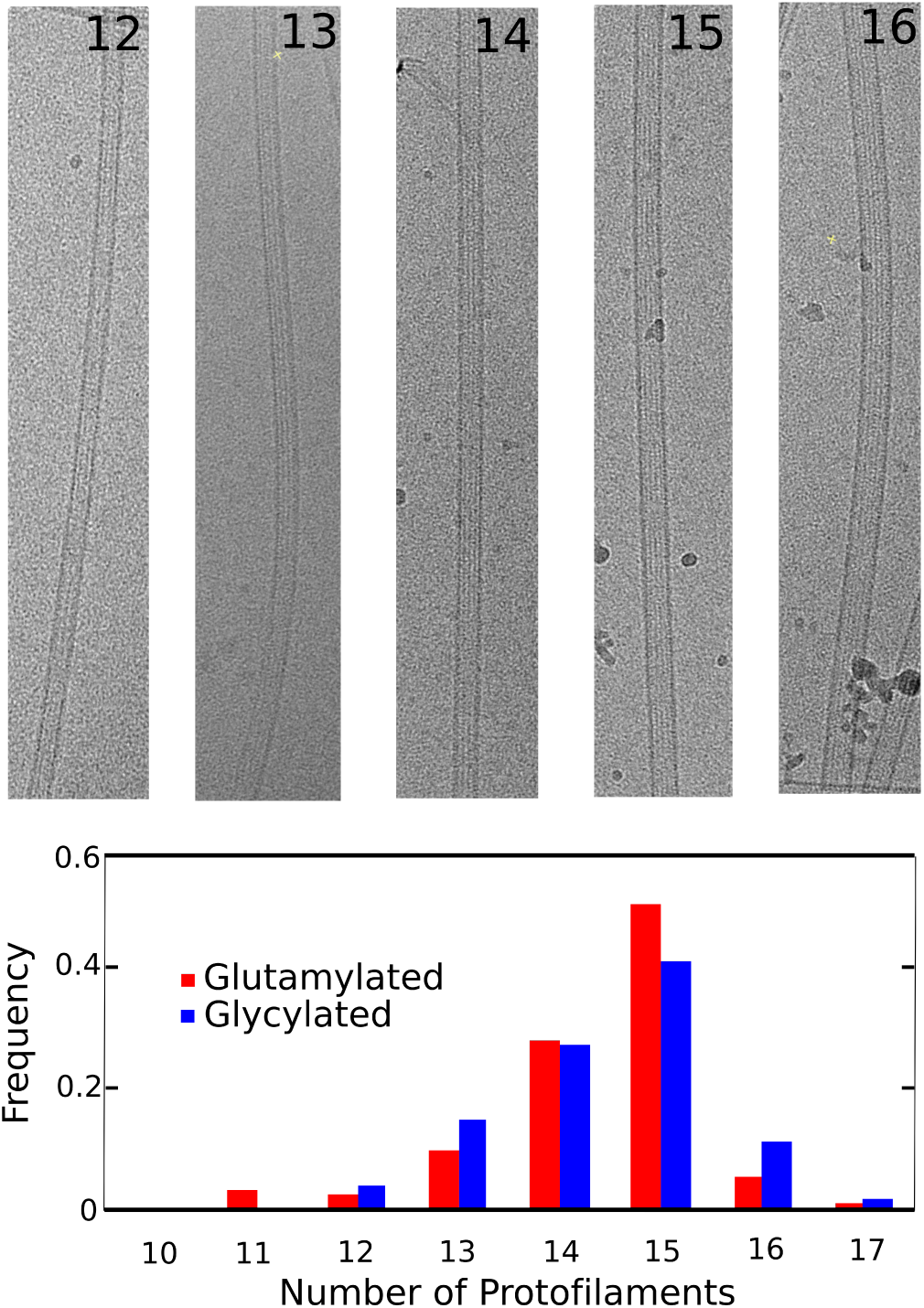
Electron microscopy representative images of varying protofilament number (top). Normalized distribution of protofilament numbers for glycylated (blue) versus glutamylated (red) microtubules (bottom). The two distributions are similar, giving a p-value of 0.97 by the K-S test.

The two differentially-modified microtubule subsets have basically indistinguishable distributions of protofilament number (K-S test p-value = 0.97). Most contain 14-15 protofilaments, as is typically seen in *in vitro* polymerization (51). We conclude that the observed differences in flexibility are due to the molecular-level interactions of the C-terminal tails rather than differences in the overall size or structure of assembled microtubules.

### Spin dynamics suggest interactions of the *α*-tubulin tail with the tubulin body

We sought to determine if NMR spectroscopy could yield insight into the mechanism by which C-terminal tail modifications affect microtubule bending rigidity. In particular, we sought to determine whether the backbone dynamics of the *α*-tubulin tail were different than that of the *β*-tubulin tail which could explain why the the bending rigidity is sensitive to the *α*-tail modifications but apparently not to the *β*-tail modifications. Difference in backbone dynamics between *α* and *β* C-terminal tails may indicate that they interact differently with the tubulin body surface.

We performed R_1_ and R_2_ NMR ^15^N-relaxation experiments to probe the local dynamics of the C-terminal tails in the presence of the tubulin dimer (52, 53). For comparison, we performed the same experiments using unmodified C-terminal tail peptides attached to GST homodimers (Fig. 6). Our low sample concentrations values coupled with typically low values for disordered domains reduced the utility of heteronuclear ^1^H-^15^N experiments, which were all below noise (NOE values <0.6). As a result, we did not attempt to apply quantitative interpretations based on combinations of these values. Fig. 6 shows the values of the R_1_ and R_2_ relaxation rates of the two C-terminal tails plotted as a function of the amino acid position relative to the typically defined start of the tails, residues D431 and D427 for *α* and *β* respectively. The shaded lines are linear fits to guide the eye. We compared the measured values for the C-terminal tails as part of the tubulin dimer in our wild type, or primarily glycylated sample, and of the C-terminal tails as unmodified peptides attached to GST dimers, to mimic their N-terminal attachment. The latter gives an indication of the intrinsic differences due to the amino acid sequence, and not to any interactions as part of the full dimer or as a result of post-translational modifications.

**Figure 6:**
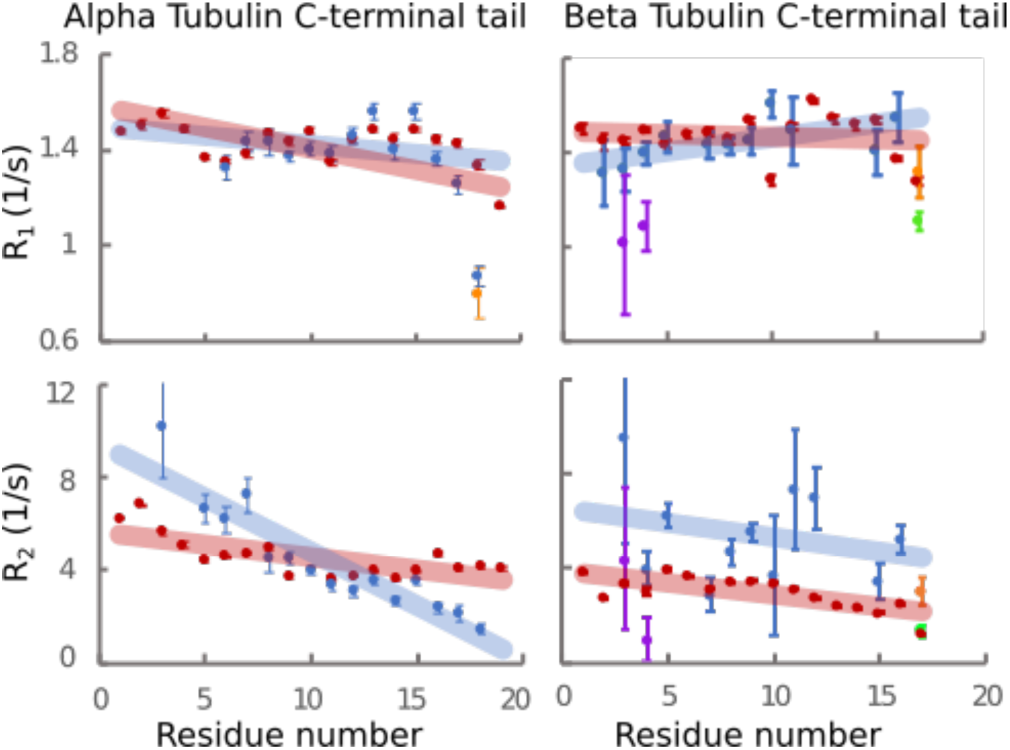
R_1_ (top row) and R_2_ (bottom row) relaxation rates for *α*- (left) and *β*- (right) tubulin tails as peptides attached to GST (red) or as part of the full tubulin dimer (blue, orange, yellow and green). Several residues in our NMR spectra are present in two different chemical environments(34). For residues near the C-termini, these correspond to modification state (orange: polyglycylation of the i-1 residue, green: mono-glycylation of the i-1 residue). For residues near the N-termini, one set of peaks is brighter (blue) and the other dimmer (purple).

We see a clear trend as a function of distance from the tubulin body in the *α*-tubulin C-terminal tail values of R_2_, which is not apparent in either R_1_ of *α* or R_1_ or R_2_ of *β* tubulin. Increased values of R_2_ are typically taken as an indication of increased interactions of the appropriate timescale, and so we conclude that the more N-terminal portions of the *α*-tubulin tail interact more significantly with the tubulin body that do residues on that tail further from the surface. Interestingly, the average value of R_2_ for the *β*-tubulin tail were intermediate, similar to those of the middle of the *α*-tubulin tail. However, the lack of clear trend along the sequence indicates that these values are dominated by local interactions, rather than interactions with the tubulin dimer surface. In contrast, the R_1_ relaxation values, typically attributed to overall motion of individual bonds is almost identical between the peptide and C-terminal tail of the *α*-tubulin tail.

Our NMR spectrum contains more than one resonance for several of the residues in the C-terminal tail. At the C-terminal extreme, we can distinguish species according to the modification state of the adjacent residue. For example, the terminal asparagine on *β*-tubulin is present in our sample in three species of differing modification state (Fig. 6, Supplementary Materials Fig. S4). The measured relaxation rates differ based on the modification state, with both R_1_ and R_2_ being larger for polyglycylated residues relative to either the unmodified (A18) or monogylcylated (B17) residues.

Residues of the tail near the tubulin body are also present in two environments, which we attributed to differences in the environment of the underlying tubulin body, although we have not been able to determine the source of these differences (34) For the *β*-tubulin tail, the segment from residues B2 to B5 (ATAE) are present in two groups corresponding to two different environments for this segment. Interestingly, the relaxation values were easy to measure for one of the groups, and are shown in blue with the rest of the chain in Fig. 6. The other group is dimmer, and more overlapped with other peaks in the spectrum. The apparent difference in relaxation rates between the two groups is further evidence of these residues having different interactions with the tubulin body.

## CONCLUSION

Post-translational modifications of tubulin dimers have been linked to both intrinsic and extrinsic properties of microtubules (6, 7). In this study, we investigated the effect of glycylation and glutamylation on the intrinsic flexural rigidity of microtubules. We purified tubulin out of three different *T. thermophila* strains, producing three pools of differentially-modified tubulin: (1) glycylated, (2) glutamylated, and (3) glycylated on only the *β*- C-terminal tail. We confirmed the extent of these modifications by mass-spectrometry and western blot. We determined that glycylation and glutamylation do not appear to change the number of protofilaments incorporated into the microtubule lattice, and therefore, do not significantly affect the size of microtubules.

Despite being of the same size, glycylated microtubules appeared significantly stiffer than both glutamylated microtubules and those lacking any post-translational modifications on the *α*-tubulin tail. The addition of high concentrations of magnesium caused the glycylated microtubules to become more flexible and more similar to that of glutamylated microtubules. Glycylated microtubules are more sensitive to changes in divalent salt concentration. This effect may be in part due to the proposed collapsed conformation of the C-terminal tails upon glycylation (33). The collapsed tail may interact more strongly with the tubulin body, an interaction which is reduced in the presence of a high concentration of magnesium.

A clear dependence of the R_2_ ^15^N-relaxation rates of the *α*-tubulin tails as a function of distance along the tail from the tubulin body surface suggests that the *α*-C-terminal tail residues interact with the tubulin body. This interaction could explain the importance of glycylation of the *α*-C-terminal tail on microtubule stiffness. The *α*-tubulin tail projects towards the adjacent dimer, supporting the view that inter-dimer rather than intra-dimer interactions may dominate the bending rigidity(54).

Glycylation on the C-terminal tails may contribute to overall stiffness and structural integrity of ciliary microtubules. Depletion of TTLL3 enzymes leads to destabilization of already assembled cilia in *T.thermophila* (27) and reduced frequency of primary cilia in mammalian cells (22, 29). In *Tetrahymena*, the TTLL3(A-F)-KO strain (lacking glycylation) has poor growth, shorter cilia, and failure of cilia to elongate in the presence of taxol (27). Glycylation is thought to be localized primarily on the B-tubules within cilia while the A-tubules are predominantly unmodified (55). The tips of cilia contain microtubules with no polyglycylation (55). While glutamylation is localized to the outer doublets in cilia and missing from the central pair (56), excessive glutamylation may destabilize and shorten cilia (57). Depletion of glycine ligases results in hyperglutamylation and vice-versa, making interpretation of any phenotype diffi cult. For example, hyperglutamylation stimulates katanin and spastin activity (17) and causes a defect in tubulin turnover (45). Extrinsic factors (such as MAPs) could contribute to ciliary stiffness (48, 58). The intrinsic differences in flexibility solely due to the modifications suggests a possible role for glycylation, especially in cellular structures undergoing high stresses such as the cilia.

Post-translational modifications play a significant role in the regulation of microtubules. We demonstrate that post-translational modifications alone regulate intrinsic flexibility of microtubules. Although the *α*- and *β*-C-terminal tails often contain similar modifications, we found that they have different effects on flexibility. This difference is most likely due to the *α*- C-terminal tail’s proximity to the dimer-dimer interface. Our results provide insight into the complex mechanism of how glycylation and glutamylation alter microtubule mechanical properties.

## AUTHOR CONTRIBUTIONS

KPW designed research, performed research, analyzed data, and wrote the manuscript. HH analyzed flexural rigidity data. TL collected and analyzed mass spectrometry data. CP collected and analyzed electron microscopy data. TLH contributed analytic tools and analyzed data for flexural rigidity. LH designed research and wrote manuscript.

## ACKNOWLEDGEMENTS

We thank Jacek Gaertig (University of Georgia) for his generous gift of *Tetrahymena thermophila* strains and polyglutamylation antibody. We thank the CU Boulder Central Analytical Mass Spectrometry Core Facility for performing LC-MS/MS experiments. This work utilized ThermoFisher LTQ orbitrap velos that was purchased with funding from W.M. Keck Foundation. The imaging work was performed at the BioFrontiers Institute Advanced Light Microscopy Core. Spinning disc confocal microscopy was performed on Nikon Ti-E microscope supported by the BioFrontiers Institute and the Howard Hughes Medical Institute. We thank Joe Dragavon (BioFrontiers Institute, University of Colorado Boulder) for his training and expertise. Electron microscopy was done at the University of Colorado, Boulder EM Services Core Facility in the MCDB Department, with the technical assistance of facility staff and Gerry Morgan (Molecular, Cellular and Developmental Biology, University of Colorado Boulder). We would also like to acknowledge J. Richard McIntosh (Molecular, Cellular and Developmental Biology, University of Colorado Boulder) for advice. The 800 MHz spectrometer was purchased and supported by the NIH (RR011969, RR16649) and the NSF (DBI-0230966, 960241).

## Supplementary Materials

**Figure S1.**
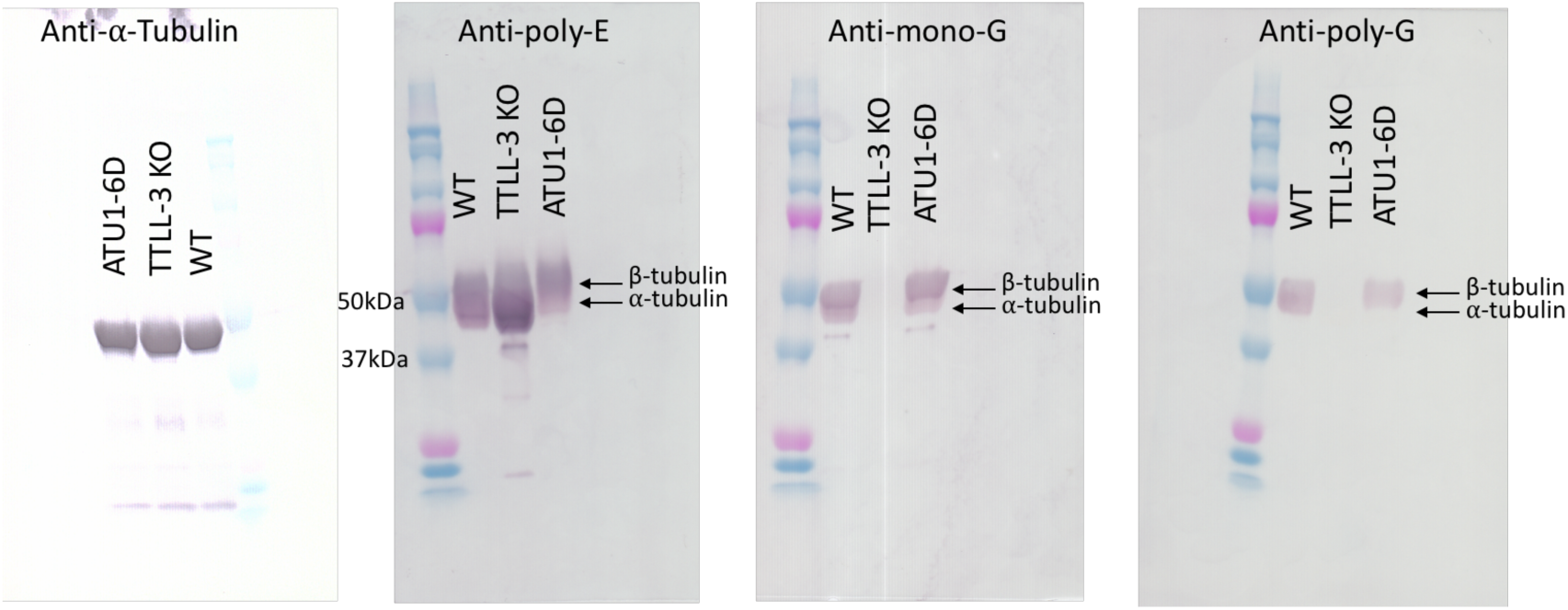
Western blot of tubulin purified from different T. thermophila strains. A qualitative western blot comparing the bulk modifications on each of the three tubulin pools purified from different strains of *T. thermophila*. The wild-type tubulin has all types of modifications detected. As expected and previously shown (Wloga et al., 2008, 2009), the TTLL3(A-F)-KO tubulin has increased glutamylation when compared to the wild-type tubulin and no apparent glycylation (either mono- or poly-). The ATU1-6D tubulin has limited modifications on the *α*-subunit. There is a low degree of poly-glutamylation and mono-glycylation on *α*- tubulin, but no poly-glycylation detected.

**Figure S2.**
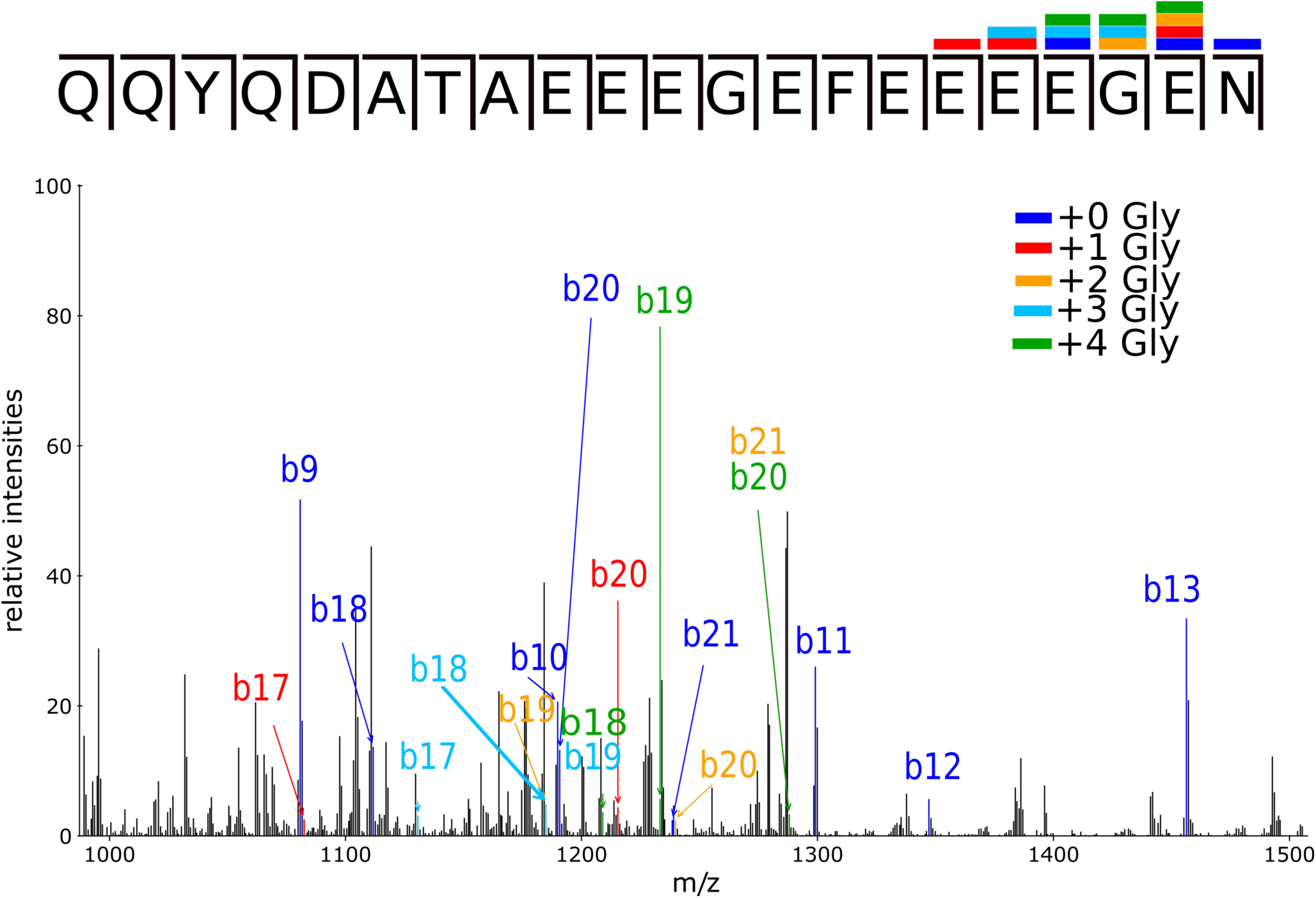
Heterogeneous MS/MS mapping of purified cell body wild-type tubulin *β*-C- terminal tail peptide. MS-MS spectra of the parent ion corresponding to mass of four additional glycine residues. The modification additions are heterogeneous with no clear single arrangement of glycine additions on this peptide. For simplicity, only the b-ions are labeled.

**Figure S3.**
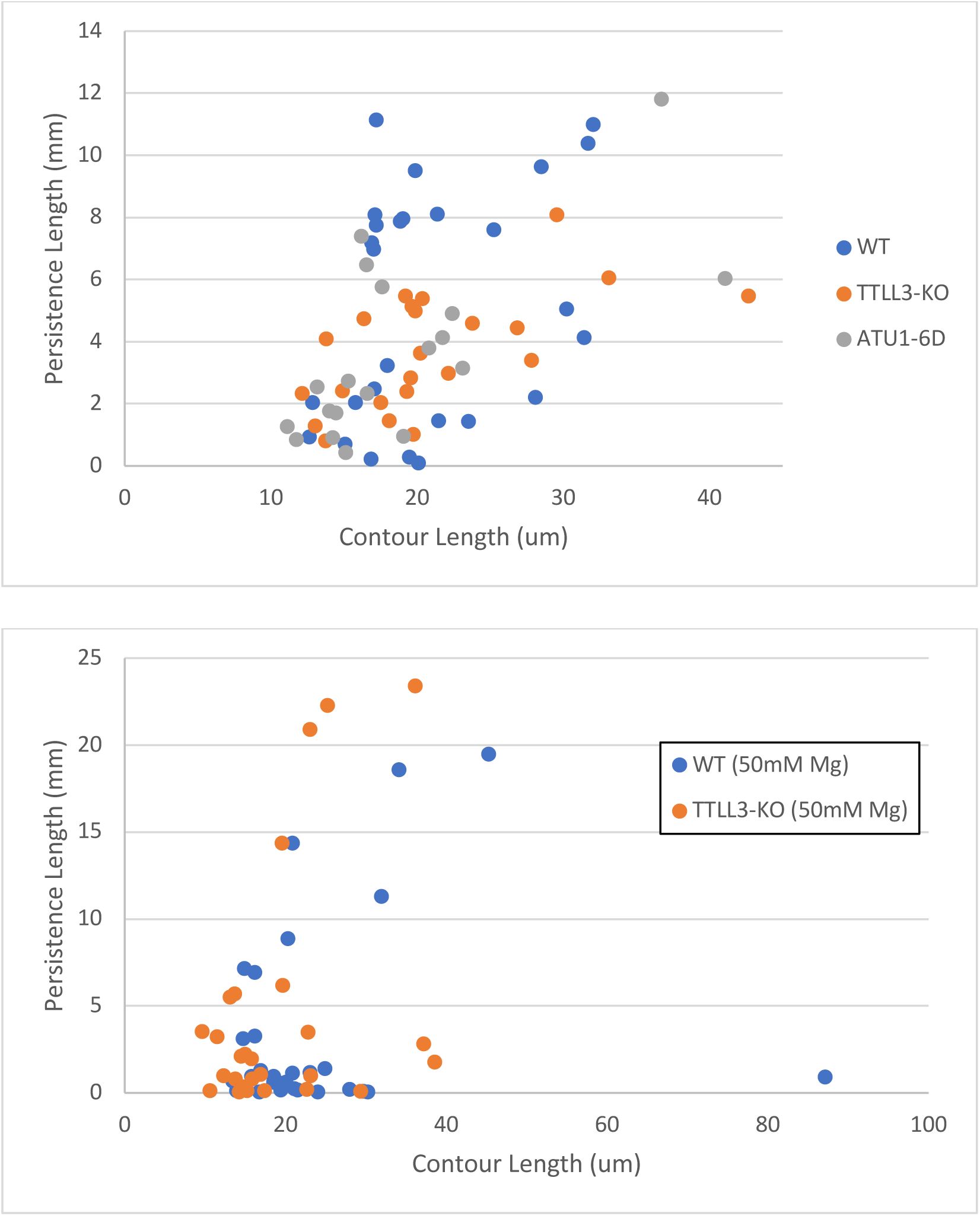
Comparison of contour length versus persistence length. (top) Scatter plot of contour length versus persistence length of microtubules in BRB80. (bottom) Scatter plot of contour length versus persistence length of microtubules in high magnesium conditions.

**Figure S4.**
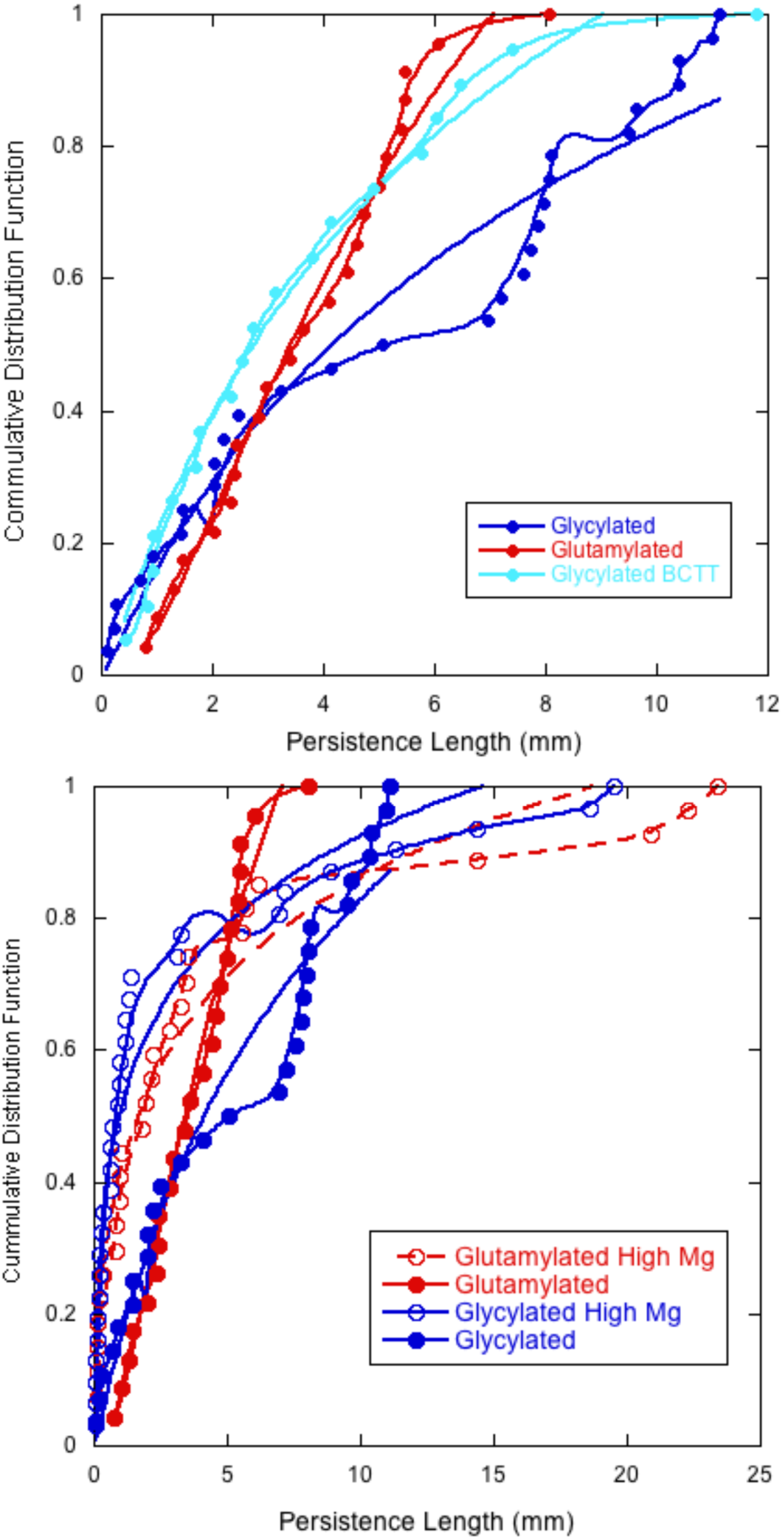
Cumulative Distribution Functions with appropriate fits. Function for CDF fit to 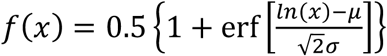. Microtubule persistence length distributions are normal or lognormal, so data is displayed as cumulative distribution functions (CDF). When compare to probability distribution function (PDF), CDFs require fewer fit parameters, produce less uncertainties in their fits, and do not involve user to binning the data.

**Figure S4.**
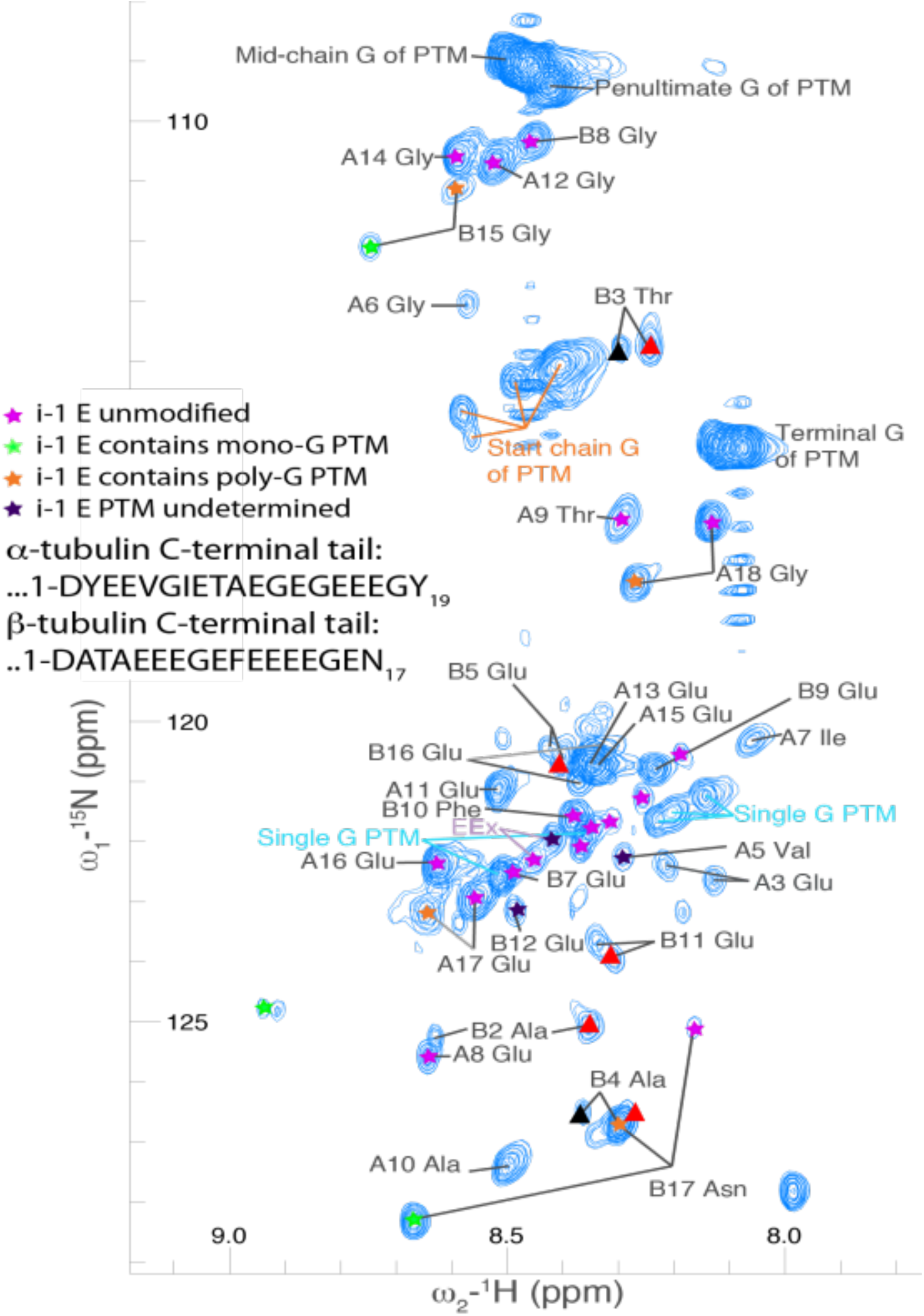
Assignment of C-terminal tails of tubulin. HNCO of C-terminal tails of tubulin annotated with the assigned peaks and corresponding modifications (Wall 2016). Added are the split chains from the beginning of the *β*-C-terminal tail (brighter ones from the relaxation plots are labeled with red triangles, dimmer are labeled with black triangles).

**Figure S5.**
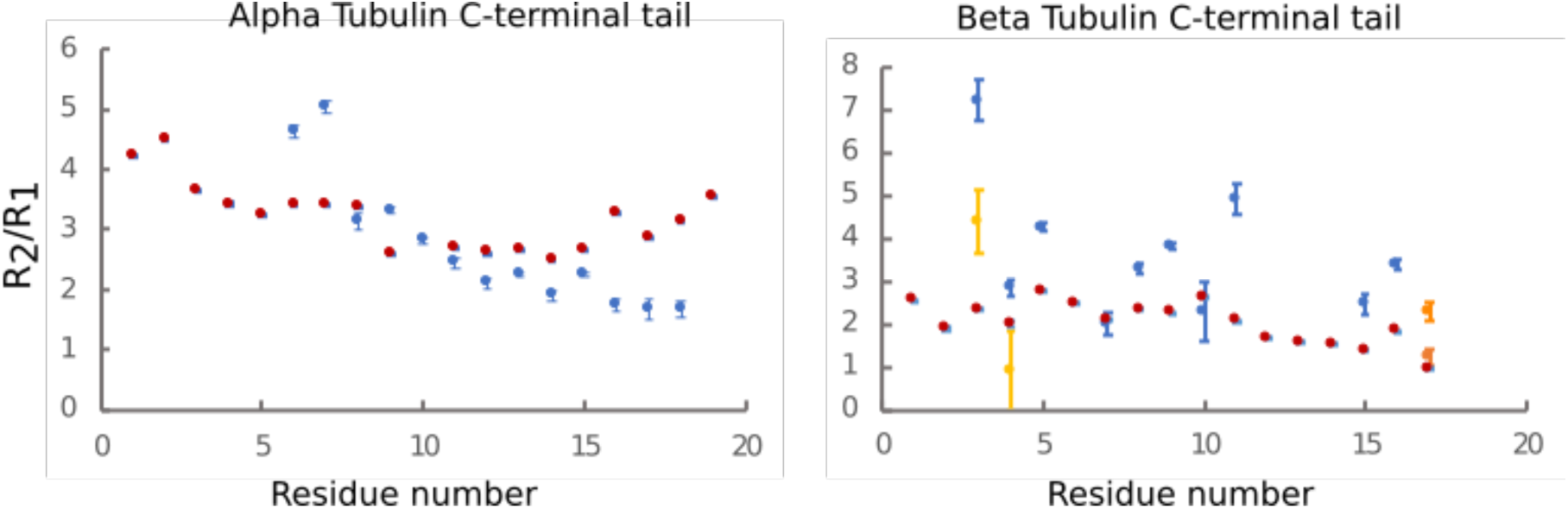
R_2_/R_1_ Relaxation. Ratios of R_2_/R_1_ plotted by residue number for the *α* and *β*-C-terminal tails.

**Table S1.** Pairwise Statistics. P-values calculated by K-S Test.

| **p-value** | TTLL3-KO<br>BRB80 | ATU1-6D BRB80 |
| --- | --- | --- |
| WT BRB80 | 0.007 | 0.042 |
| TTLL3-KO BRB80 | xxx | 0.731 |

| **p-value** | WT (BRB80) | TTLL3-KO<br>(BRB80) | WT (50mM Mg) |
| --- | --- | --- | --- |
| WT (50mM Mg) | <0.001 | <0.001 | xxx |
| TTLL3-KO Mg | 0.050 | 0.045 | 0.223 |

Reference: http://www.physics.csbsju.edu/stats/KS-test.html

**Table S2:** Persistence length and fit parameters for cumulative distribution fit. Function for CDF fit to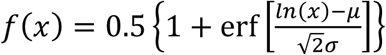.

| | $n$ | $L_p$ (mm) | $\mu$ | $\sigma$ | $R^2$ |
| --- | --- | --- | --- | --- | --- |
| Glycylated | 28 | $3.24 \pm 1.60$ | $2.72 \pm 0.07$ | $1.92 \pm 0.13$ | 0.95 |
| Glutamylated | 23 | $0.64 \pm 0.18$ | $1.96 \pm 0.03$ | $1.07 \pm 0.05$ | 0.98 |
| Glycylated β-CTT | 19 | $0.55 \pm 0.42$ | $2.20 \pm 0.04$ | $1.76 \pm 0.07$ | 0.98 |
| Glycylated (50mM Mg) | 31 | $0.90 \pm 0.30$ | $2.68 \pm 0.14$ | $4.01 \pm 0.22$ | 0.94 |
| Glutamylated (50mM Mg) | 27 | $1.49 \pm 0.37$ | $2.93 \pm 0.01$ | $3.57 \pm 0.17$ | 0.96 |

